# Yap1-driven intestinal repair is controlled by group 3 innate lymphoid cells

**DOI:** 10.1101/769190

**Authors:** Mónica Romera-Hernández, Patricia Aparicio-Domingo, Natalie Papazian, Julien J. Karrich, Ferry Cornelissen, Remco M. Hoogenboezem, Janneke N. Samsom, Tom Cupedo

## Abstract

Tissue repair requires temporal control of progenitor cell proliferation and differentiation to replenish damaged cells. In response to acute insult, group 3 innate lymphoid cells (ILC3) regulate intestinal stem cell maintenance and subsequent tissue repair. ILC3-derived IL-22 is important for stem cell protection, but the mechanisms of ILC3-driven tissue regeneration remain incompletely defined. Here we report that group 3 innate lymphoid cell (ILC3)-driven epithelial proliferation and tissue regeneration are independent of IL-22. In contrast, ILC3 amplify the magnitude of Hippo-Yap1 signaling in intestinal crypt cells, ensuring adequate initiation of tissue repair and preventing excessive pathology. Mechanistically, ILC3-driven tissue repair is Stat3-independent, but involves activation of Src family kinases. Our findings reveal that ILC3-driven intestinal repair entails distinct transcriptional networks to control stem cell maintenance and epithelial regeneration which implies that tissue repair and crypt proliferation can be influenced by targeting innate immune cells independent of the well-established effects of IL-22.

---

The intestinal epithelial lining forms a physical barrier that prevents translocation of commensal microorganisms and defects in intestinal barrier integrity or maintenance have severe clinical impact (Konig et al., 2016; Peterson and Artis, 2014). Loss of barrier function activates deleterious immune responses and the ensuing mucositis can lead to transmural ulcers and the need for parenteral nutrition (Sonis, 2004). Such complications are a major dose-limiting side effect of high-dose chemotherapy for head-and-neck cancers or during myeloablative conditioning for hematopoietic stem cell transplantation as part of leukemia treatment (Keefe, 2007; Sonis, 2004). Insufficient intestinal barrier repair is also a pathological feature underlying inflammatory bowel disease (IBD), including Crohn’s disease and ulcerative colitis (Hollander et al., 1986; Odenwald and Turner, 2013). Reduced barrier integrity evokes continuous bacterial translocation, fueling reactivation of microbiota-specific T cells and subsequent disease recurrence in up to 40% of IBD patients in remission (Munkholm et al., 1994). Enhancing intestinal epithelial repair, also coined mucosal healing, has become a sought-after result in experimental and clinical IBD research, yet the cells and signals that enhance epithelial regeneration are still ill defined (Dulai et al., 2015; Florholmen, 2015; Neurath, 2014; Shah et al., 2016).

In contrast to the potentially deleterious consequences of overt anti-bacterial immune activation following barrier loss, innate immune cell-derived signals are also implicated as positive regulators of intestinal regeneration (Aparicio-Domingo et al., 2015; Lindemans et al., 2015). Group 3 innate lymphoid cells (ILC3s) safeguard epithelial stem cells after acute small intestinal damage inflicted by Methotrexate, mediated through production of IL-22 (Aparicio-Domingo et al., 2015). IL-22 is constitutively produced by small intestinal ILC3 (Savage et al., 2017), is important for homeostatic production of antimicrobial peptides (Liang et al., 2006), and drives stem cell proliferation in organoid cultures ex-vivo (Lindemans et al., 2015).

Intestinal repair is driven by epithelial stem cell differentiation, and subsequent proliferation of progenitor cells in crypts of LieberkÜhn (Barker, 2014). Crypt regeneration is regulated by highly conserved signaling pathways activated by signals from local niche cells (Kabiri et al., 2014; Medema and Vermeulen, 2011; Sato et al., 2011). Wnt and Notch signals control stem cell self-renewal (Clevers et al., 2014), while in addition, Wnt favors Paneth cell differentiation (Van Es et al., 2005), while Notch controls the absorptive versus secretory lineage choice(Jones et al., 2015). Hedgehog signaling is involved in progenitor proliferation (Van Dop et al., 2010), while BMPs activate the pericryptal mesenchyme to support the epithelial stem cell niche(He et al., 2004). The Hippo-YAP1 pathway is mostly active in response to damage, inhibiting Wnt and Notch signaling to drive early differentiation (Gregorieff et al., 2015; Imajo et al., 2015). IL-22 was postulated to also be a stem cell niche factor, derived not from structural cell but from intestinal ILC3, and involved in intestinal repair (Hanash et al., 2012; Pickert et al., 2009).

In this study, we set out to define the role of ILC3-derived IL-22 during epithelial repair and to mechanistically interrogate this process. We show that in response to acute small intestinal damage the stem cell protective cytokine IL-22 is dispensable for crypt cell proliferation and that in contrast, ILC3s modulate intestinal repair by controlling the amplitude of Hippo-YAP1 signaling. Yap1 activation involves signals downstream of gp130 and Src family kinases. These findings unveil a unique layer of epithelial regulation, in which evolutionary conserved regenerative pathways are amplified by cells of the innate immune system. This realization paves the way for novel therapeutic strategies aimed at enhancing tissue repair during IBD or during cytotoxic anti-cancer therapies through activation of intestinal immune cells independent of the well-established effects of IL-22 (Hanash et al., 2012; Lindemans et al., 2015).

## RESULTS AND DISCUSSION

### Crypt cell proliferation after stem cell damage is IL-22 independent

The ILC3-derived cytokine IL-22 positively impacts the percentage of Lgr5^+^ intestinal stem cells that can be recovered from small intestinal crypts after acute damage, and induces epithelial proliferation in vitro (Aparicio-Domingo et al., 2015; Lindemans et al., 2015). To determine the importance of IL-22 for small intestinal regeneration we compared intestinal pathology and crypt cell proliferation after MTX-induced damage in IL-22 deficient mice and ILC3-deficient ROR*γ*t^−/−^ mice. As we have shown before (Aparicio-Domingo et al., 2015), absence of ILC3 increased tissue pathology in response to MTX. In contrast, pathology in IL-22 deficient mice was similar to pathology of IL22-sufficient littermate controls **(Figure 1A)**. As a measure of crypt regeneration, we enumerated proliferating crypt cells during the regenerative phase of the response (day 4). This revealed that the severely reduced proliferation, characteristic of ILC3-deficiency (Aparicio-Domingo et al., 2015), was absent from IL-22^−/−^ animals, in which crypt proliferation was indistinguishable from littermate controls **(Figure 1B)**. This led us to hypothesize that small intestinal crypt proliferation after acute MTX-induced damage can occur in an IL-22 independent manner, uncoupled from IL-22 dependent epithelial stem cell maintenance.

**Figure 1.**
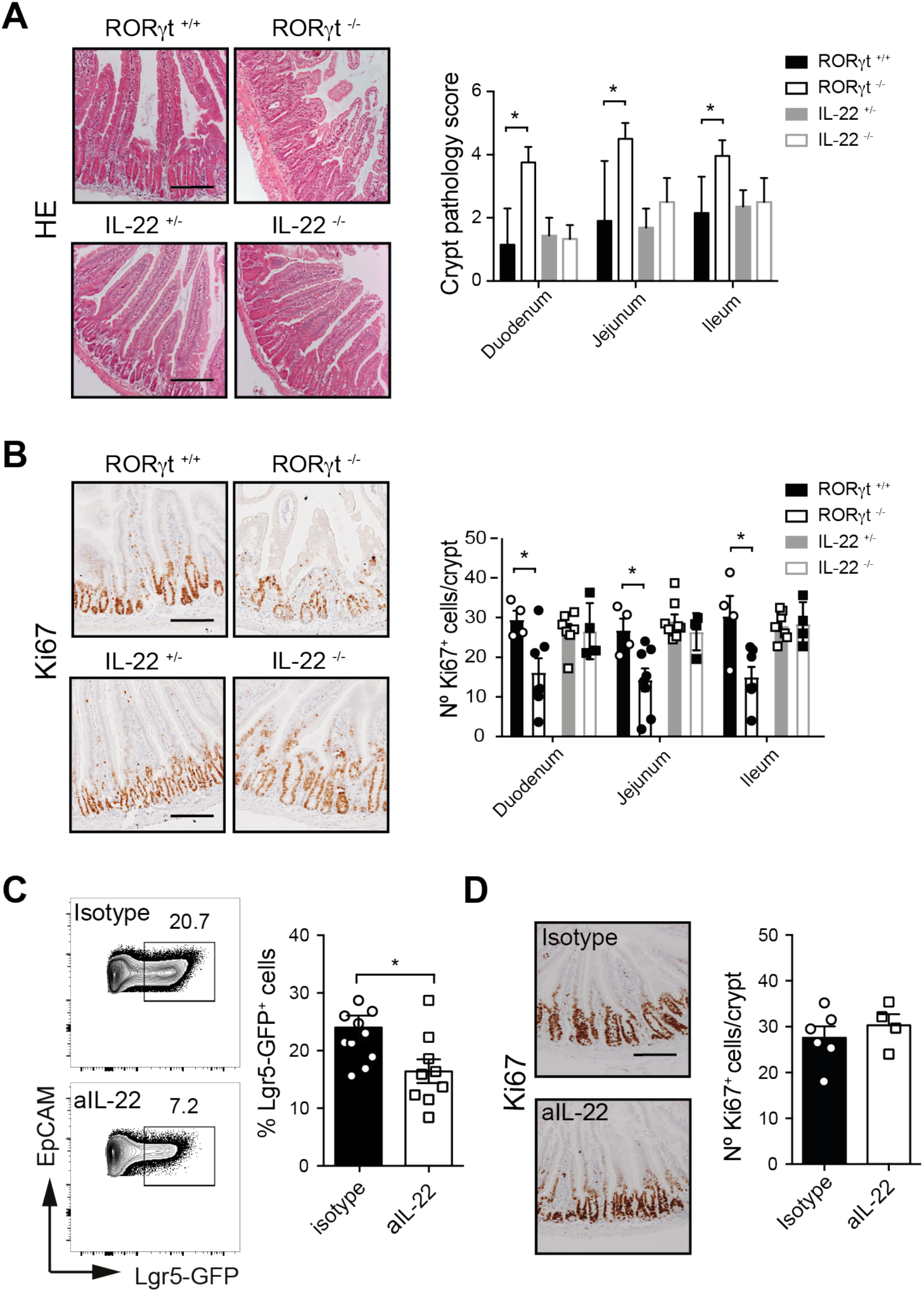
Differential regulation of crypt proliferation and stem cell maintenance. **(A)** Representative H&E staining and crypt histopathology scores of small intestinal sections from ROR*γ*t^−/−^, IL-22^−/−^ and littermate control mice four days after MTX treatment. **(B)** Representative Ki67 immunostaining and number of Ki67^+^ cells in small intestinal crypts of ROR*γ*t^−/−^, IL-22^−/−^ and littermate control mice four days after MTX treatment. **(C)** Representative flow cytometry plots and frequency of Lgr5-GFP^+^ cells in duodenal crypts four days after MTX from Lgr5-GFP^+/−^ mice treated with neutralizing anti-IL-22 antibodies. Numbers adjacent to outlined areas indicate percentage of Lgr5-GFP^+^ cells within Live(+)CD45(-)Ter119(-)CD31(-)EpCAM1(+) cells. **(D)** Representative Ki67 immunostaining and number of Ki67^+^ cells in duodenal crypts four days after MTX in Lgr5-GFP^+/−^ mice treated with neutralizing anti-IL-22 antibodies or isotype controls. Unpaired Mann-Whitney test *p<0.01; **p<0.001; statistically not significant (not indicated). Scale bars: 50μm; n=4-8 mice per group.

To test this hypothesis directly, we neutralized IL-22 by antibody treatment in Lgr5-GFP stem cell-reporter mice during exposure to MTX. This allowed for combined analyses of stem cell maintenance and crypt proliferation. Inhibition of IL-22 signaling was confirmed by downregulation of transcripts encoding the prototypic IL-22 target genes *Reg3g* and *Reg3b* **(Figure S1)**. Upon neutralization of IL-22 signaling with antibodies **(Figure 1C)** during tissue damage, maintenance of Lgr5-GFP-expressing cells was impaired. In contrast, crypt proliferation was unaffected **(Figure 1D)** and was similar to control animals. Together these data reveal that ILC3-dependent stem cell maintenance and crypt proliferation are mechanistically distinct responses to insult, and that the early proliferative response following small intestinal injury is IL-22 independent.

### YAP1 signaling drives small intestinal regeneration

Given that small intestinal crypt proliferation occurred in an IL-22 independent manner, we set out to identify the epithelial repair programs activated by acute small intestinal damage. To this end, we analyzed changes in the epithelial stem cell transcriptome evoked by MTX immediately following induction of damage (day 1) and during regeneration (day 4). RNA-sequencing of purified Lgr5-GFP^hi^ crypt cells revealed rapid transcriptional induction of target genes of the regeneration-associated Hippo/YAP1 pathway, at one and four days after damage **(Figure 2A).** Activation of YAP1 involves protein de-phosphorylation and subsequent nuclear translocation. Under homeostatic conditions, YAP1 localization in villus epithelium was exclusively cytoplasmic, while in crypt epithelial cells, both cytoplasmic and faint nuclear localization were observed, suggestive of continuous low grade YAP1 activation **(Figure 2B)**. MTX exposure induced YAP1 nuclear translocation in crypt epithelial cells **(Figure 2B)**, in line with the increased transcription of YAP1 target genes. Concomitant to YAP1 activation, transcription of genes indicative of WNT **(Figure 2C)** and Notch **(Figure 2D)** signaling were reduced, in agreement with YAP1-dependent temporal repression of stem cell self-renewal, favoring rapid differentiation at the expense of stemness (Barry et al., 2013; Gregorieff et al., 2015; Yui et al., 2018). Indeed, transcription of genes associated with stem cell identity were reduced **(Figure 2E)**, while transcription of genes associated with mature secretory cells, including goblet cells (GC), Paneth cells (PC) and enteroendocrine cells (EEC) were all increased **(Figure 2F and Figure S2A-C)**, as was the proportion of phenotypic enteroendocrine cells in isolated crypts **(Figure 2G and Figure S2D)**.

**Figure 2.**
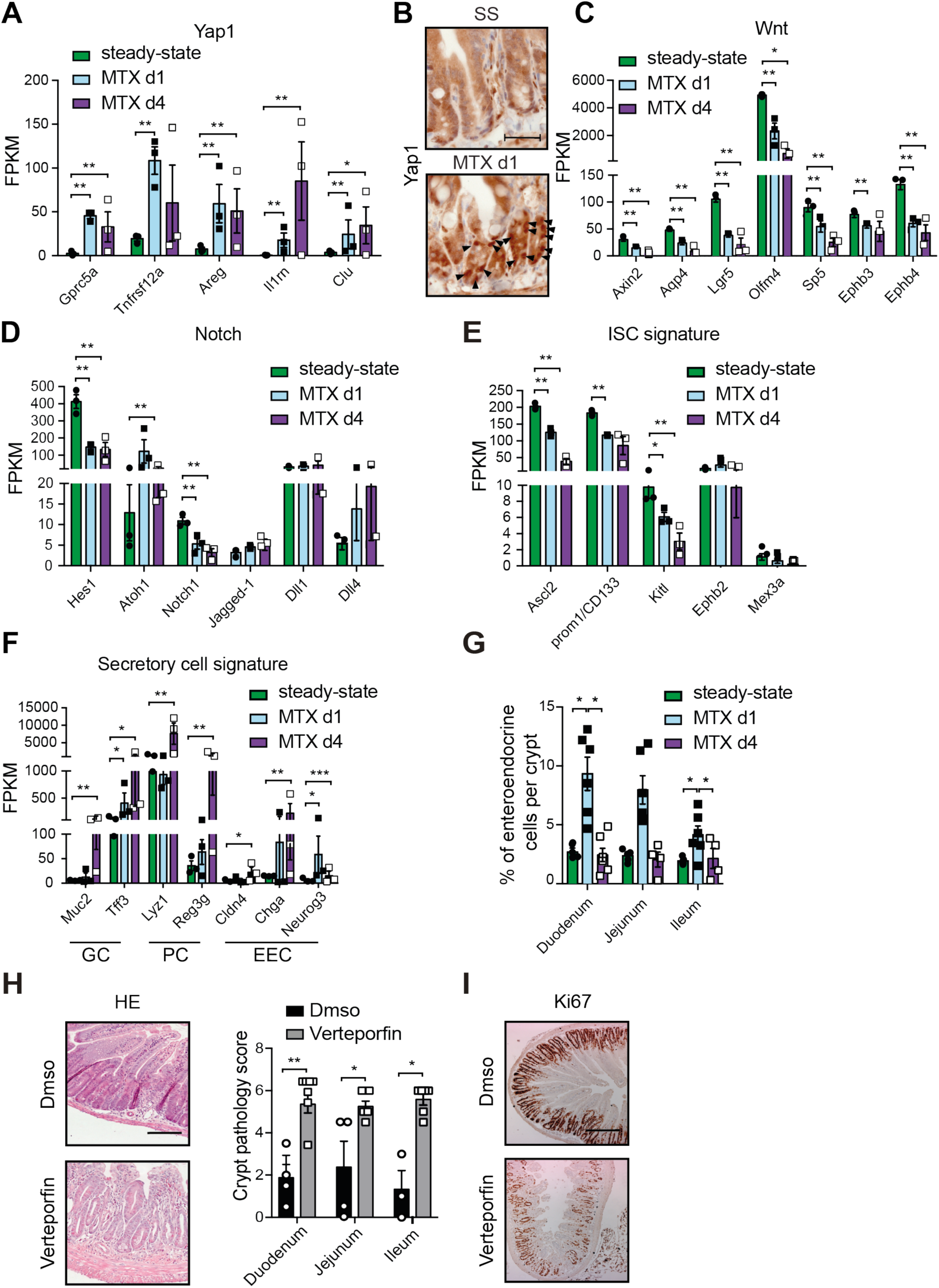
YAP1 activation drives acute small intestinal regenerationy. **(A)** FPKM values of Yap1 target genes in LGR5-GFP^hi^ crypt cells analyzed by RNA-sequencing at steady-state (SS), one and four days after MTX. **(B)** Representative YAP1 immunostaining of duodenal sections at the indicated time points after MTX exposure. FPKM values derived from RNA-sequencing of LGR5-GFP^hi^ crypt cells of genes involved in **(C)** Wnt-signaling, **(D)** Notch-signaling, **(E)** intestinal stem cell (ISC) signature and **(F)** secretory cell identity, including goblet cells (GC), Paneth cells (PC) and enteroendocrine cells (EEC). **(G)** Percentages of EEC in small intestinal crypt-derived cell suspensions analyzed by flow cytometry at the indicated time points. EEC were gated as Live(+)CD45(-)Ter119(-)CD31(-)EpCAM1(+)Lgr5-GFP(-)CD24^hi^SSC^lo^ cells. **(H)** Representative H&E staining and crypt pathology scores of small intestine four days after MTX exposure in the presence of Verteporfin or DMSO control. **(I)** Representative Ki67 immunostaining of duodenal sections four days after MTX treatment in the presence of Verteporfin or DMSO control. FPKM values are plotted for transcripts with statistically significant Log_2_fold change (DESeq2 analysis of count data) with ** adjusted p value <0,01, * adjusted p value <0.05 or statistically not significant (not indicated). Unpaired Mann-Whitney test **p<0.01; *p<0.05 (G, H). Scale bars: 20μm (B), 50 μm (H), 100 μm (I). n=3 mice per group (A, C-G); n=4-8 mice per group (B, H, I).

YAP1 activates target gene transcription by binding to nuclear TEAD transcription factors (Zhao et al., 2008). The interaction between YAP1 and TEADs can be inhibited by Verteporfin, an FDA-approved drug used for treatment of macular degeneration (Battaglia Parodi et al., 2016). To resolve the importance of YAP1 activation for small intestinal regeneration after MTX-induced damage we blocked YAP1-TEAD interactions during MTX exposure. YAP1 inhibition increased tissue pathology, characterized by severe crypt abnormalities **(Figure 2H)** and a loss of crypt cell proliferation **(Figure 2I)**, very similar to the loss of regeneration in ILC3-deficient ROR*γ*t^−/−^ mice. Together these findings identify YAP1 activation as a dominant driver of early small intestinal crypt regeneration following acute damage.

### YAP1 activation is blunted in the absence of ILC3

Based on the importance of YAP1 activation and ILC3 presence for small intestinal regeneration we hypothesized that activation of the Hippo/YAP1 pathway is controlled by ILC3 presence. To test this hypothesis, we analyzed YAP1 activation in mice lacking ILC3. In ROR*γ*t-deficient mice, damage-induced YAP1 nuclear translocation was strongly reduced **(Figure 3A)**, leading to a decrease in the percentage of crypts containing at least one cell per transverse section with nuclear YAP1 **(Figure 3B)**, and thus an increase in crypts failing to activate YAP1. To molecularly define crypt responses in the absence of ILC3 we crossed Lgr5-GFP reporter mice to ILC3-deficient ROR*γ*t^−/−^ mice and analyzed the epithelial stem cell transcriptome after small intestinal damage by RNA-sequencing. YAP1 activation in LGR5-GFP^hi^ epithelial cells from mice lacking ILC3 was severely blunted, and relative to homeostasis, no significant changes could be detected in the vast majority of YAP1 target genes **(Figure 3C)**. Diminished YAP1 activation was concomitant with failure to reduce WNT **(Figure 3D)** and Notch **(Figure 3E)** target genes and signaling components, and an absence of both loss of stemness **(Figure 3F)** and induction of transcripts involved in mature secretory cell differentiation **(Figure 3G)**. Similarly, the transient increase in enteroendocrine cell differentiation seen in control crypts **(Figure 2G)** was completely absent in crypts from ROR*γ*t^−^/ ^−^ mice **(Figure 3H)**, highlighting the functional consequences of reduced YAP1 activation. Together these data reveal that the magnitude of the evolutionary conserved, regeneration-associated, Hippo-YAP1 pathway is amplified by presence of small intestinal ILC3.

**Figure 3.**
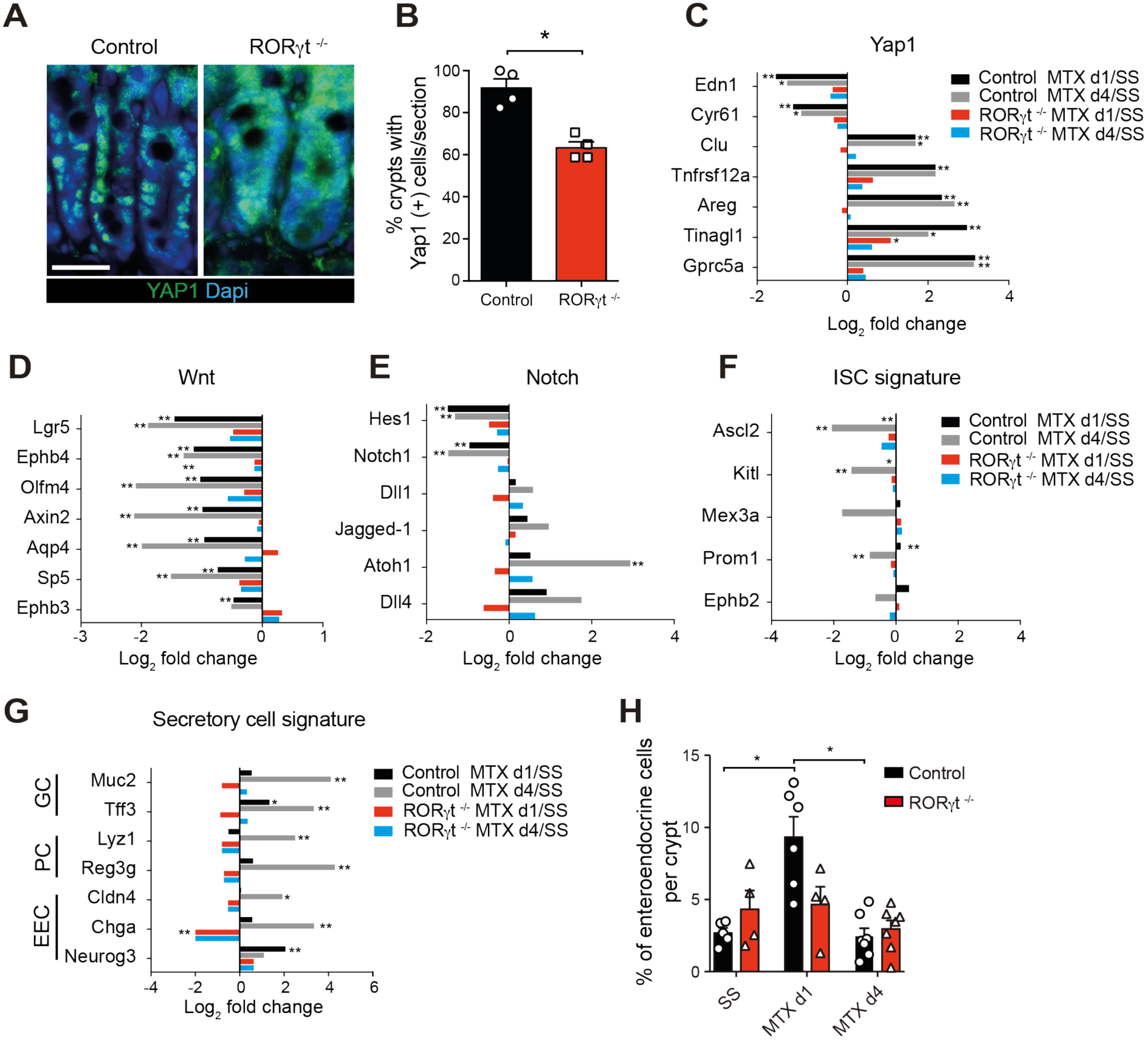
YAP1-activation is blunted in the absence of ILC3. **(A)** Representative YAP1 immunostaining of duodenal crypts from ROR*γ*t^−/−^ mice and littermate controls one day after MTX. Scale bar 50 μm. **(B)** Percentage of duodenal crypts containing at least 1 cell with nuclear translocation of YAP1 in ROR*γ*t^−/−^ mice and littermate controls at one day after MTX. **(C-G)** Log_2_ fold change values determined by RNA-sequencing of Lgr5-GFP^hi^ crypt cells comparing MTX d1 versus SS (steady state) for control mice (black bars) or ROR*γ*t^−/−^ mice (red bars) and MTX d4 versus SS for control mice (grey bars) or ROR*γ*t^−/−^ mice (blue bars) showing **(C)** YAP1 target genes, **(D)** Wnt and **(E)** Notch related genes, **(F)** intestinal stem cell genes and **(G)** secretory cell genes. **(H)** Percentages of enteroendocrine cells determined by flow cytometry in duodenal crypts at the indicated time points from ROR*γ*t^−/−^ mice (red bars). Control mice (as in Figure 2G) shown for comparison (black bars). Enteroendocrine cells were gated as Live(+)CD45(-)Ter119(-)CD31(-)EpCAM1(+)Lgr5-GFP(-)CD24^hi^SSC^lo^ cells. Log_2_ fold change (DESeq2 analysis of count data) with ** adjusted p value <0,01, * adjusted p value <0.05 or statistically non-significant (not indicated) (C-F). FPKM values are plotted for transcripts that have statistically significant Log_2_fold change (DESeq2 analysis of count data) with ** adjusted p value <0,01, * adjusted p value <0.05 or statistically non-significant (not indicated). Unpaired Mann-Whitney test *p<0.01; statistically not significant (not indicated) (B,H). n=4-8 mice per group (A,B,H). n=3 mice per group (C-G).

### Activation of YAP1 signaling in intestinal crypts is independent of IL-22

Our findings indicate that crypt proliferation and Lgr5 cell maintenance are independently regulated, fueled by YAP1 and IL-22 respectively. To assess whether IL-22 has a role in epithelial YAP1 activation after small intestinal damage we quantified YAP1 activation in epithelial stem cells from mice exposed to MTX in the presence of IL-22 neutralizing antibodies. Lgr5-GFP^hi^ crypt cells were analyzed by RNA sequencing one day after MTX. The efficacy of IL-22 neutralization was controlled by analysis of Reg3*γ* and Reg3β transcription in purified Paneth cells **(Figure S3A).** IL-22 neutralization did not alter transcripts associated with YAP1 **(Figure S3B**, WNT **(Figure S3C)** or Notch **(Figure S3D)** targets or signaling components, nor transcripts involved in stemness **(Figure S3E)** or secretory cell differentiation **(Figures S3E-F)**. These data illustrate that IL-22 signaling is dispensable for YAP1 activation following acute damage to the small intestine.

### YAP1 activation requires gp130 dimerization and Src family kinase activation

Damage-induced epithelial YAP1 activation can occur in response to a variety of stimuli, including signaling through the IL-6 family receptor gp130 and mechanotension in response to altered extracellular matrix composition (Taniguchi et al., 2015). Mechanotension-induced YAP1 activation occurs during the later stages of tissue repair and involves activation of a discrete set of genes(Yui et al., 2018). Analyses of these mechanotension-associated genes in Lgr5-GFP^hi^ stem cells after MTX-induced damage showed that these genes are either not transcribed or transcribed at similar levels in ILC3-deficient and control mice **(Figure S4A-B)**, making it unlikely that this pathway is important during the initial ILC3-depemdent activation of epithelial YAP1.

In order to define whether ILC3-driven YAP1 involved gp130 signaling we first analyzed the transcriptome of Lgr5-GFP^hi^ small intestinal crypt cells for presence of gp130 and the cytokine-specific receptor chains that multimerize with gp130 to form functional receptor units. Under homeostatic conditions, as well as after damage, Lgr5-GFP^hi^ crypt cells transcribed gp130, the il11ra1 chain, and low levels of Cntfr **(Figure 4G)**. IL-6 can also signal via a soluble IL-6R that complexes with gp130, thereby precluding the necessity for cell-autonomous IL-6R transcription (Novick et al., 1989). The IL-6 and IL-11 pathways have been associated with multiple aspects of tissue regeneration, and alterations in these pathways are linked to IBD susceptibility and intestinal cancer progression (Garbers and Scheller, 2013; Katsanos and Papadakis, 2017; Taniguchi and Karin, 2014). The IL-6 and IL-11 receptors utilize homo-dimerization of gp130 to form a specific receptor with a single IL-6R or IL-11R chain. To analyze possible involvement in Yap1 activation of gp130 homo-dimerization we used a small molecule inhibitor of this process (LMT-28) (Hong et al., 2015). LMT-28 application during the induction of small intestinal damage with MTX reduced nuclear translocation of YAP1 in crypt epithelial cells, reduced the percentage of crypts containing at least one cell with nuclear YAP1 per transverse section, and thus increased the number of crypts that failed to activate YAP1-driven regeneration **(Figure 4H)**. Downstream of gp130-associated receptors signals are transduced by either STAT3 signaling or by activation of Src family kinases (SFKs) (Taniguchi et al., 2015). To assess the importance of these two pathways we inhibited STAT2/3 signaling with the small molecule inhibitor STATTIC and SFK activity with the pharmacological inhibitor PP2. Efficiency of STAT inhibition was controlled by analyses of known STAT3 target genes **(Figure S4C)**. Inhibition of STAT3 signaling also blocks IL-22 receptor signaling highlighted by the failure of Lgr5^+^ stem cell maintenance following MTX damage **(Figure 4C)**. In contrast, the YAP1-dependent proliferative response at day 4 post MTX was not affected by STAT2/3 inhibition **(Figure 4D)** indicating that STAT molecules are not involved in this process. Administration of the SFK inhibitor PP2 prevented nuclear translocation of YAP1 in small intestinal crypt cells **(Figure 4E)** and resulted in increased crypt pathology and loss of crypt proliferation **(Figure 4F)**, reminiscent of the regenerative defects in ILC3-deficient ROR*γ*t^−/−^ mice. These findings again highlight the dichotomy between stem cell maintenance, which is both ILC3 and IL22-dependent, and crypt proliferation that is ILC3-dependent yet IL22-independent.

**Figure 4.**
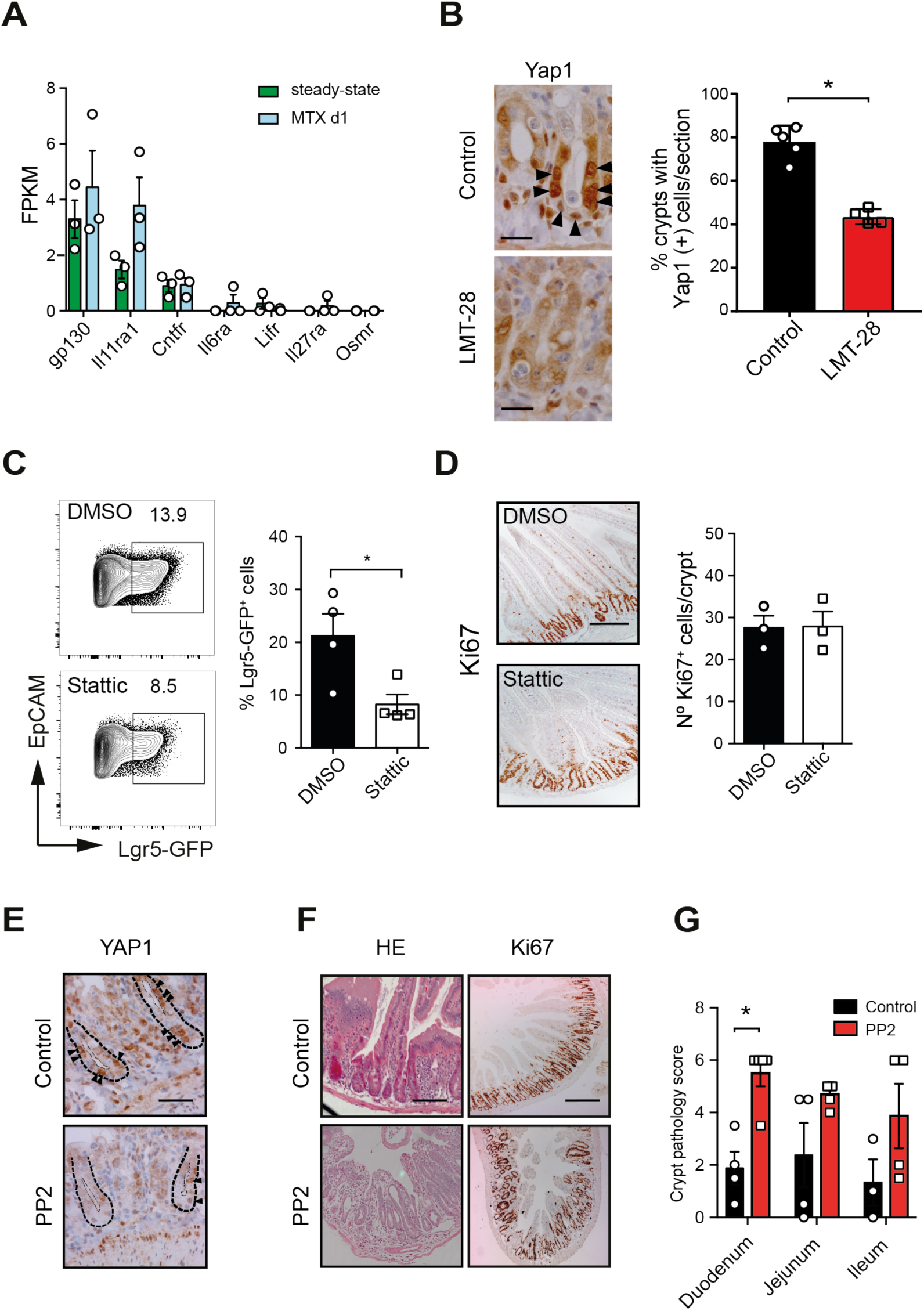
YAP1 activation involves gp130 – SFK signaling. **(A)** FPKM values of gp130 and gp130 co-receptors expressed by Lgr5-GFP^hi^ crypt cells at steady-state and one day after MTX. **(B)** Representative YAP1 immunostaining and percentage of duodenal crypts containing at least 1 cell with nuclear translocation of YAP1 at one day after MTX treatment in the presence of LMT-28 or vehicle control. **(C)** Representative flow cytometry plots and frequency of Lgr5-GFP^+^ cells in duodenal crypts four days after MTX from Lgr5-GFP^+/−^ mice treated with STATTIC or DMSO vehicle control. Numbers adjacent to outlined areas indicate percentage of Lgr5-GFP^+^ cells within Live(+)CD45(-)Ter119(-)CD31(-)EpCAM1(+) cells. **(D)** Representative Ki67 immunostaining and number of Ki67^+^ cells in duodenal crypts four days after MTX in Lgr5-GFP^+/−^ mice treated with STATTIC or DMSO vehicle control. **(E)** Representative YAP1 immunostaining in duodenal crypts from mice treated with SFK-inhibitor PP2 or DMSO control at one day after MTX. **(F)** Representative H&E and Ki67 staining of duodenal sections four days after MTX treatment in the presence of PP2 or DMSO control. **(G)** Crypt pathology score of small intestinal sections four days after MTX in mice treated with PP2 or DMSO control. FPKM values are plotted for transcripts that have statistically significant Log_2_fold change (DESeq2 analysis of count data). Unpaired Mann-Whitney test *p<0.01 (B), unpaired T test *p<0.05 (C,D). Scale bar 100μm (D, F) and 50 μm (B). n=3 mice per group (A, C, D) n=4-6 mice per group (B, E-G).

Collectively, our findings reveal that the evolutionary conserved, regeneration-associated Hipo-YAP1 signaling pathway is amplified by presence of ILC3, independent of the stem cell-protective effect of IL-22. This implies that immune cell-driven tissue regeneration is the sum of multiple parallel pathways, including the well-established IL22R signaling pathway as well as ILC3-dependent YAP1 activation as shown in this report. The mechanistic uncoupling of stem cell protection and crypt regeneration could suggest that ILC3 act on not only on LGR5-expressing intestinal progenitors, but also on additional damage-associated epithelial precursor subsets (Ayyaz et al., 2019; Tian et al., 2011).

YAP1-driven intestinal regeneration is critical for epithelial proliferation after intestinal damage in insects (Karpowicz et al., 2010; Ren et al., 2010; Shaw et al., 2010). Nevertheless, ILC3 are present only in mammals and fish (Hernandez et al., 2018), and are lacking from more primitive organisms (Lane et al., 2009). Combined with our findings, this implies that evolution favored a complementary layer of crypt regulation driven by specialized innate immune cells. This paves the way for future design of novel targeting strategies aimed at activating ILC3-driven tissue repair in IBD or reduce side effects of anti-cancer therapies, independent of IL-22.

## ACKNOWLEDGMENTS

Neutralizing IL-22 antibodies and IL22-deficient mice were kindly provided by Genentech. This work was supported by ZonMW Innovational Research Incentives Vidi grant #91710377 to T. Cupedo, Veni grant #91615128 to FC, and by the People Program (Marie Curie Actions) of the European Union’s Seventh Framework Program FP7/2007-2013 under REA grant agreement no. 289720.

## Author Contributions

Conceptualization, TC and JNS; Investigation, MRH PAD NP JJK and FC; Formal analysis, MRH PAD NP JJK FC and RMH; Data curation, RMH; Writing – Original draft MRH and TC; Writing – Review and Editing, TC and JNS; Supervision, TC and JNS; Funding Acquisition, TC and FC.

## METHODS

### Mice

C57BL/6, B6.129P2(Cg)-Rorc^tm2Litt^/J (ROR*γ*t-GFP), IL-22^−/−^ (Kindly provided by Genentech, South San Francisco, CA), Lgr5-GFP-IRES-creERT2 (Lgr5-GFP) and Lgr5-GFP/ROR*γ*t-GFP mice were bred at the animal facility of the Erasmus University Medical Center Rotterdam. Animal experiments were approved by relevant authorities and procedures were performed in accordance with institutional guidelines. Age and gender-matched littermates were used whenever possible.

### Small intestinal damage

8-12 weeks old mice were injected i.p. with 120 mg/kg clinical grade Methotrexate PCH at day-1 and with 60 mg/kg at day 0. Tissues were collected at day 1 and day 4 after the last MTX injection.

### Cytokine modulation during MTX treatment

150 μg anti-IL-22 antibody (8E11, kindly provided by Genentech, South San Francisco, CA) or mouse IgG1 isotype control (MOPC-21, BioXCell) were administered i.p to Lgr5-GFP^+/−^ mice every 2 days, starting 4 days before the first MTX dose, until day 2 after the last MTX dose. Stattic (Sigma) was injected at day −1 and day 0 at 0.5 mg/ml in 2.5% DMSO. Control mice were injected with 2.5% DMSO. Verteporfin (Selleckchem) and PP2 (Sigma) were administered at day −1, day 0 and day 1 and mice were analyzed at day 4. Verteporfin was dissolved at a final concentration of 10 mg/injection and PP2 at 1 mg/injection. Both were dissolved in 10%DMSO. Control mice received 10% DMSO. LMT-28 was injected i.p at 5 mg/kg, at days −1, 0 and 2. Control mice were given the same volume of saline containing 10% ethanol.

### Crypt isolation

Isolation of intestinal crypts was performed as previously described (Sato et al., 2009). Briefly, isolated small intestines were opened longitudinally and washed with cold PBS. 5 mm pieces were washed with cold PBS, incubated in EDTA (2 mM) at 4°C for 30 min and resuspended in PBS. Crypt-enriched sediments were passed through a 70 μm cell strainer and centrifuged at 200 g for 2 min to separate the crypts from single cells. Crypts were incubated with 1 ml of TrypLE Express (Gibco) + Dnase I (Merk Millipore) at 37°C for 10-15 min until crypt dissociation was observed. Single cell suspensions were filtered through 40 μm cell strainer and labeled with conjugated antibodies.

### Antibodies

The following antibodies were used for flow cytometry: CD45 (30F11; Invitrogen); EpCAM-1 (G8.8), CD24 (M1/69), CD31 (390), Ter119 (TER-119;); all from BioLegend. Dead cells were excluded with 7AAD (Beckman coulter). Antibodies used for paraffin immunostaining were: rat anti-mouse Ki67 monoclonal antibody (MIB-5, Dako) and rabbit anti-mouse Yap1 antibody (D8H1X, Cell signaling).

### Flow cytometry and cell sorting

Fc receptors were blocked with appropriate sera or rat-anti-mouse CD16/CD32 (2.4G2; BD Biosciences). All labelings were performed in PBS containing 2% heat-inactivated fetal calf serum (FCS) at 4°C. Labelled cells were analyzed on a FACS LSRII (BD Biosciences) and data processed with FlowJo software (FlowJo, LLC).

### RNA sequencing

cDNA was prepared using SMARTer Ultra Low RNA kit (Clontech Laboratories) for Illumina Sequencing following the manufacturer’s protocol. The Agilent 2100 Bio-analyzer and the High Sensitivity DNA kit were applied to determine the quantity and quality of the cDNA production. Amplified cDNA was further processed according to TruSeq Sample Preparation v.2 Guide (Illumina) and paired end-sequenced (2×75bp) on the HiSeq 2500 (Illumina). Demultiplexing was performed using CASAVA software (Illumina) and the adaptor sequences were trimmed with Cutadapt (http://code.google.com/p/cutadapt/). Alignments against the mouse genome (mm10) and analysis of differential expressed genes were performed with DESeq2 in the R environment on the raw fragment counts extracted from the BAM files by HTSeq-count(Gröschel et al., 2014). Cufflinks software was used to calculate the number of fragments per kilobase of exon per million fragments mapped (FPKM) for each gene.

### Histology

Small intestinal tissue pieces (5 mm) or Swiss rolls were fixed in 4% PFA (4h, room temperature), washed in 70% ethanol and embedded in paraffin. Four-μm sections were deparaffinized and stained with hematoxylin (Vector Laboratories) and eosin (Sigma-Aldrich). For Ki67 and YAP1 detection, endogenous peroxidases were blocked in 1% periodic acid in deionized water for 20 min, and antigen retrieval was achieved by microwave treatment in citrate buffer (10mM, pH 6.0). Prior to staining, Fc receptors were blocked in 10% normal mouse serum and 10% of normal serum matching the host species of the secondary antibody, 10 mM Tris buffer, 5 mM EDTA, 0.15 M NaCl, 0.25% gelatin, and 0.05% Tween-20 (pH 8.0). Tissue sections were incubated overnight at 4°C with primary antibodies in PBS supplemented with 2% normal mouse serum. Immunoreactions were detected using biotinylated donkey anti-rat (Dako) and goat-anti-rabbit (Vector Laboratories) and incubated with the Vectastain ABC Elite Kit (Vector Laboratories) and 3,3’-diaminobenzidine tetrahydrochloride (Sigma-Aldrich). Sections were counterstained with hematoxylin. Pathology scores and enumeration of Ki67 and Yap1 were performed blinded. Pathology was scored as previously described (de Koning *et al.*, 2006) and Ki67-expressing cells were counted in 7 to 15 crypts per section.

### Immunohistochemistry

Tissues were frozen in Tissue-Tek O.C.T compound (Sakura Finetek Europe B.V.) and stored at −80 °C. Six μm cryosections were fixed for 5 min in ice-cold acetone and air-dried for 10 min, and subsequently blocked with 5% normal mouse serum and 5% normal donkey serum for 15 min. Sections were incubated with primary antibody for 1 hr. at room temperature, followed by a 30min incubation with secondary antibodies. Sections were embedded in Pro-long Gold with DAPI (Invitrogen) and analyzed on a Leica DMRXA.

### Transcript analysis

RNA was extracted using the NucleoSpin RNA XS kit (Machery Nagel). RNA from sorted cells was amplified with the Ovation PicoSL WTA System V2 (NuGen) according to manufacturer’s protocol. For quantitative PCR, a Neviti Thermal Cycler (Applied Biosystems) and SensiFAST SYBR Lo-Rox kit (BioLine) were used, with the addition of MgCl_2_ to a final concentration of 4 mM. All reactions were performed in duplicate and normalized to the expression of *Gapdh or Cyclophilin*. Relative expression was calculated by the cycling threshold (CT) method as 2^−ΔCT^. The primers sequences can be found in Table 1 (see below).

**Table 1.**
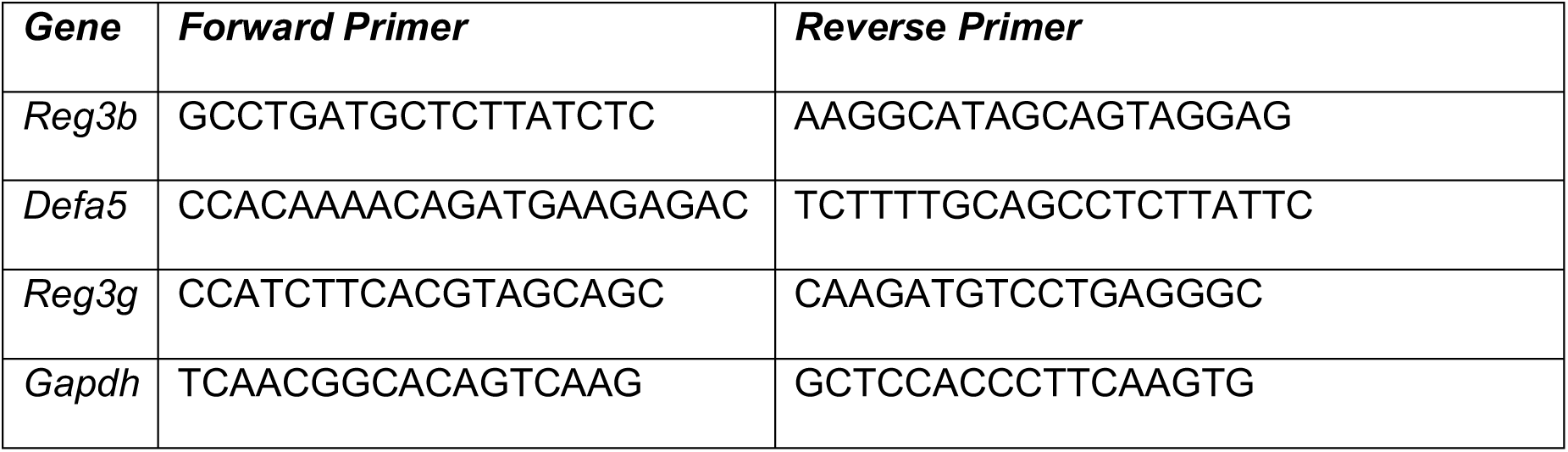
Primer sequences.

### Statistical analysis

Samples were analyzed using unpaired Mann-Whitney test or Unpaired T test as indicated in figure legends. P values <0.05 were considered significant. Data are shown as mean ± SEM.

### Data Availability

The accession number for the RNA sequencing data reported in this study is ArrayExpress: E-MTAB-6639.

