## Supplemental Figures 1-4 for "Yap1-driven intestinal repair is controlled by group 3 innate lymphoid cells"

**Control of tissue repair by innate lymphoid cell-driven amplification of epithelial Hippo-Yap1 signaling**

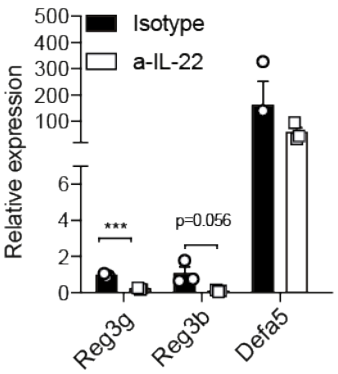


**Supplemental Figure 1: Control for antibody-mediated IL-22 neutralization.** *Reg3$\gamma$*, *Reg3$\beta$* and *Def$\alpha$5* mRNA transcript expression relative to *Gapdh* in total crypts from Lgr5-GFP$^{+/-}$ mice after isotype control or anti-IL-22 antibodies at four days after MTX. Unpaired Mann-Whitney test: ***p<0.001; statistically not significant (not indicated). n=3-5 mice per group.

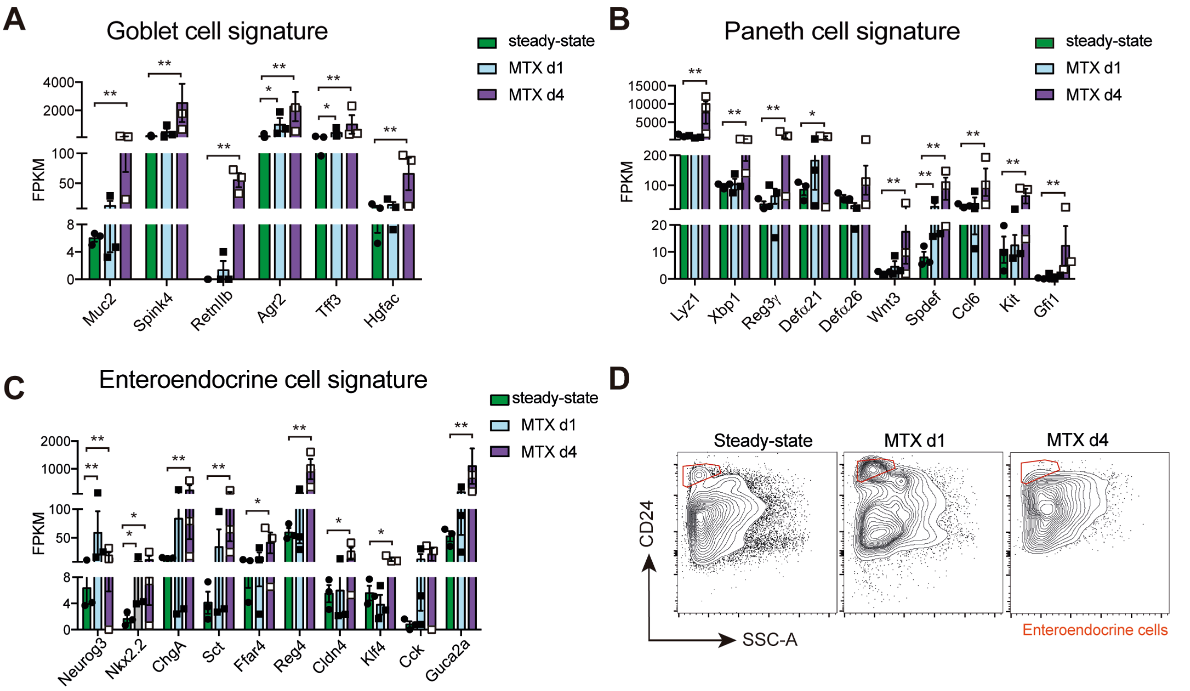


**Supplemental Figure 2. Increase of secretory cell genes and EE cells in response to MTX.** FPKM values at steady-state (SS), one and four days after MTX of genes associated with **(A)** mature goblet cells, **(B)** Paneth cells and **(C)** enteroendocrine cells. **(D)** Representative flow cytometry plots of small intestinal crypt-derived cell suspensions at the indicated time points. Indicated CD24$^{hi}$ SSC-A$^{lo}$-expressing cells correspond to enteroendocrine cells that were gated as Live cells CD45(-)Ter119(-)CD31(-)EpCAM(+)Lgr5-GFP(-)CD24$^{hi}$SSC-A$^{lo}$ cells. FPKM values are plotted for transcripts that have statistically significant Log$_2$fold change (DESeq2 analysis of count data) with ** adjusted p value <0,01, * adjusted p value <0.05 or statistically not significant (not indicated). n=3 mice per group (A-C), n=3-6 mice per group (D).

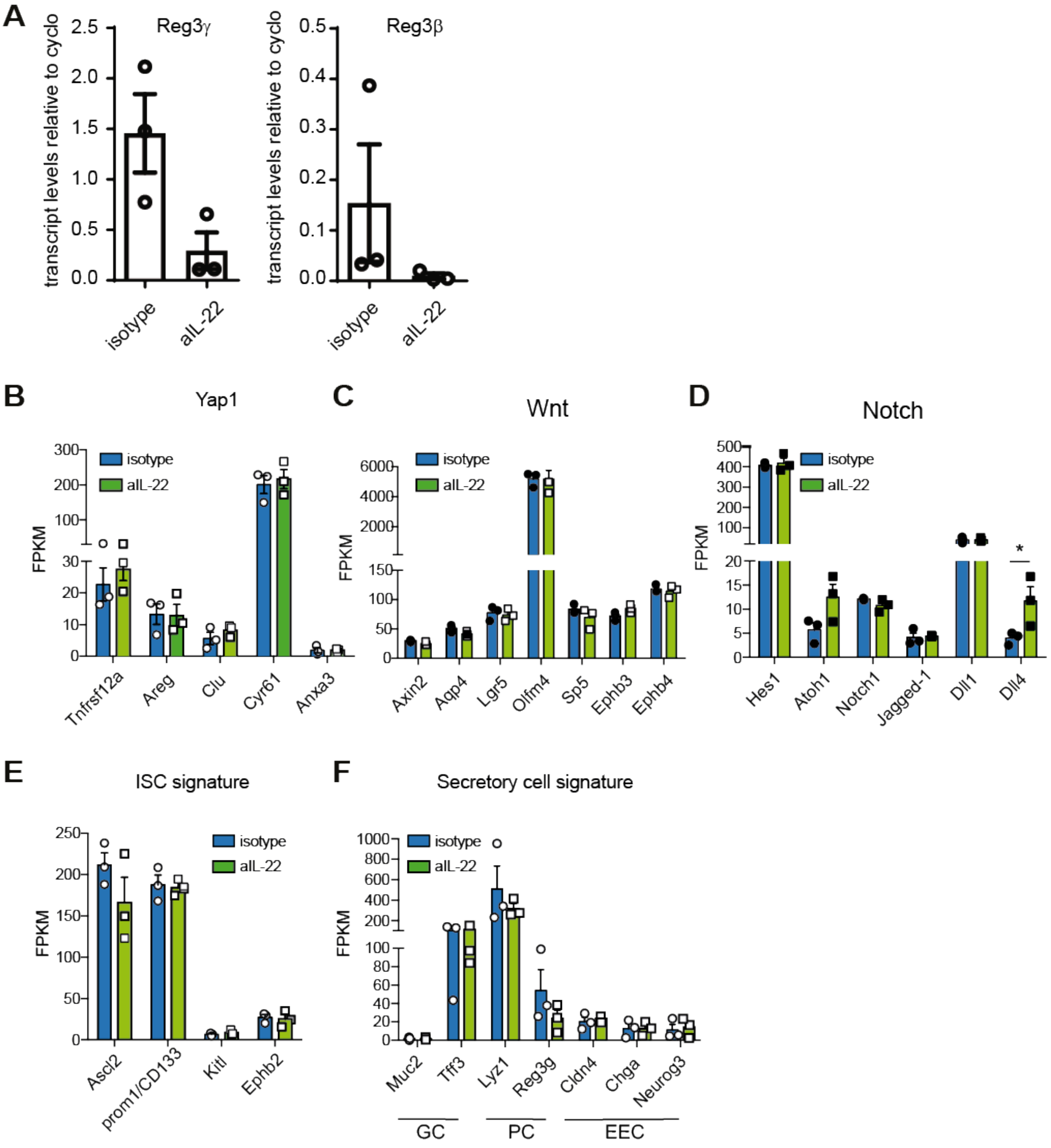


**Supplemental Figure 3. IL-22 is dispensable for Yap1 activation in Lgr5-GFP$^{hi}$ cells.**

**(A)** Transcript analyses by qPCR of FACS-sorted Paneth cells 1 day after MTX application in the presence of neutralizing IL-22 antibody or isotype control. **(B-F)** FPKM values from RNA-sequencing analyses of Lgr5-GFP$^{hi}$ crypt cells at MTX day 1 treated with neutralizing IL22 antibodies or isotype control. **(B)** YAP1 target genes and **(C)** genes associated with WNT and **(D)** Notch signaling. **(E)** ISC signature genes and **(F)** secretory cell genes including goblet cells (GC), Paneth cells (PC) and enteroendocrine cells (EEC). FPKM values are plotted for transcripts that have statistically significant $Log_2$fold change (DESeq2 analysis of count data) with ** adjusted p value <0,01, * adjusted p value <0.05 or not significant (not indicated). n=3 mice per group.

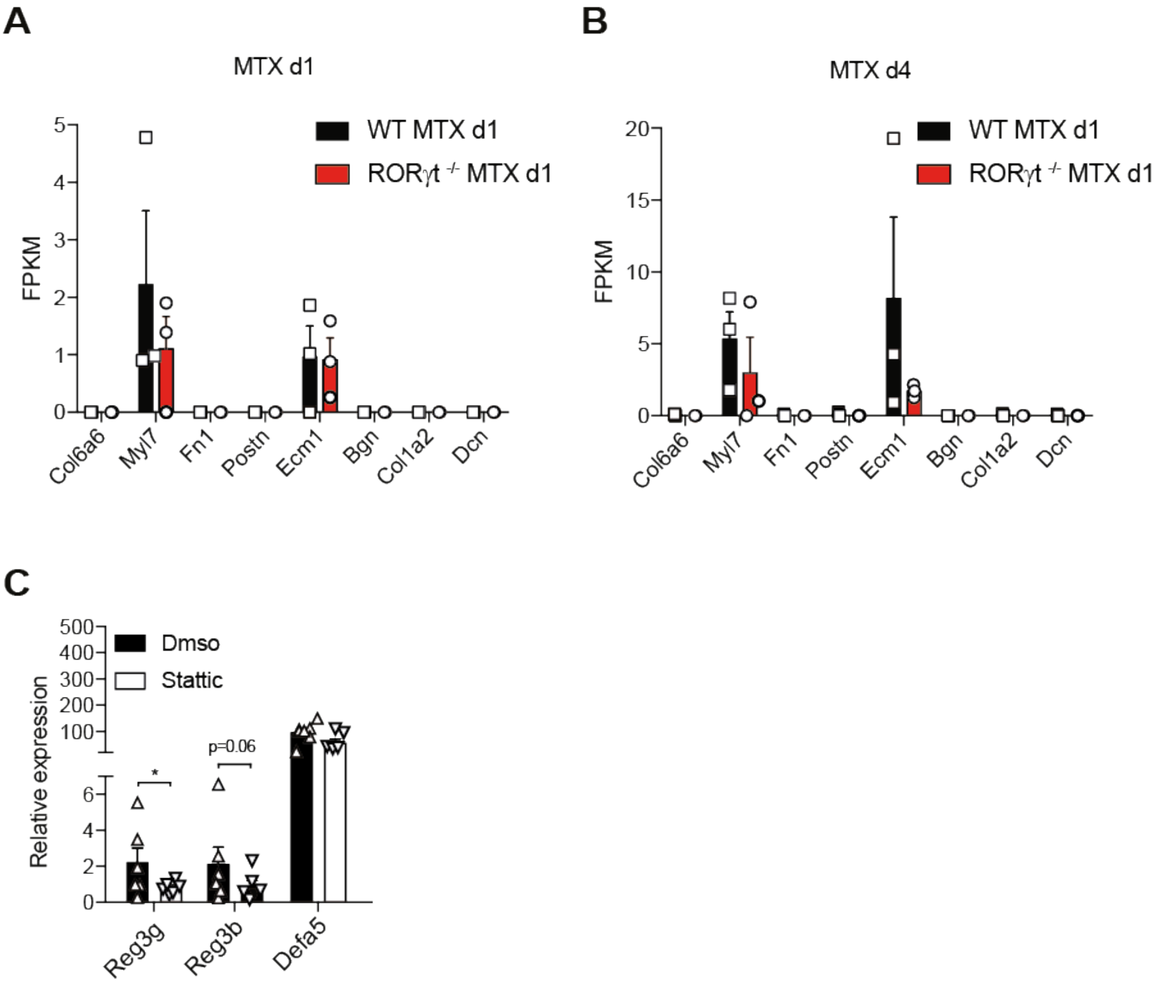


**Supplemental Figure 4. Gene signatures related to YAP1 activation.** FPKM values from RNA-sequencing of Lgr5-GFP$^{hi}$ crypt cells for genes involved in mechanotension-induced repairing epithelium at day 1 **(A)** and 4 **(B)** after MTX. Genes according to Yui et al. (Yui et al., 2018). **(C)** *Reg3γ*, *Reg3β* and *Defα5* mRNA transcript expression relative to *Gapdh* in total crypts from Lgr5-GFP$^{+/-}$ mice after STATTIC or DMSO vehicle control treatment, at four days after MTX. Unpaired Mann-Whitney test: *p<0.05; statistically not significant (not indicated). n=3-5 mice per group.
